# Advancing Spectrally-Resolved Single Molecule Localization Microscopy using Deep Learning

**DOI:** 10.1101/2022.07.29.502097

**Authors:** Hanna Manko, Yves Mély, Julien Godet

## Abstract

Spectrally-Resolved Single Molecule Localization Microscopy (srSMLM) is a recent multidimensional technique enriching single molecule localization imaging by the simultaneous recording of single emitters spectra. As for SMLM, the localization precision is fundamentally limited by the number of photons collected per emitters. But srSMLM is more impacted because splitting the emission light from single emitters into a spatial and a spectral channel further reduces the number of photons available for each channel and impairs both spatial and spectral precision - or forces the sacrifice of one or the other. Here, we explored the potential of deep learning to overcome this limitation. We report srUnet - a Unet-based image processing that enhances the spectral and spatial signals and compensates for the signal loss inherent in recording the spectral component. We showed that localization and spectral precision of low-emitting species remain as good as those obtained with a high photons budget together with improving the fraction of localizations whose signal is both spatially and spectrally interpretable. srUnet is able to deal with spectral shift and its application to multicolour imaging in biological sample is straight-forward.

srUnet advances spectrally resolved single molecule localization microscopy to achieve performance close to conventional SMLM without complicating its use.

## Introduction

Super-resolution microscopy has opened exciting new opportunities in life science to study biological processes at the nanometer scale. Among the super-resolution imaging methods, single molecule localization microscopy (SMLM) (1–3) is widely used - presumably because it builds on conventional wide-field imaging and is relatively easy to implement (4). SMLM is based on the principle that the localization of a molecule can be determined with high accuracy from its PSF if the latter corresponds to a single molecule emitter. The elegant idea behind SMLM is to achieve the spatial isolation of single emitters by repeatedly recording sparse subsets of individual fluorophores stochastically-activated over time. Isolated emitters are then pinpointed and their localizations accumulated to reconstruct a SMLM image. The routine lateral resolution of SMLM is in the range of 10-50 nm - an order of magnitude improvement over conventional diffraction-limited microscopy resolutions. SMLM has recently been extended beyond the spatial positions of single emitters to explore additional dimensions of their emission signals. In Spectrally-resolved SMLM (srSMLM), both the full emission spectra and the localizations of single emitters are simultaneously recorded (5–8). The additional characterization of the spectral component is a breakthrough and srSMLM has rapidly found successful applications in multicolour imaging (5, 8–10) or tracking (9, 11–13), single molecule conformational changes (14–17), or single-molecule polarity sensing (6, 18–20).

Different technical approaches have been reported to capture the spectral decomposition of light (21). A dual-objective configuration was initially described (5) where one objective lens is dedicated to localization measurements and the second - combined with a prism - is used for collecting spectral-measurements of the same single molecules. While this design achieved excellent light collection efficiency, more recent works reported the use of a single-objective combined either with a transmission grating (Fig. 1 a.) or with the association of a beam splitter and a prism. These setups are simpler in design and impose fewer constraints on sample mounting as compared to a dual-objective setup. But splitting the signal into two paths reduces the number of photons available for each path. Additionally, strong spectral dispersion reduces the signal-to-noise ratio in the spatial domain, so that parameters of the emission spectra cannot always be recovered or properly characterized for all detected molecules particularly for the lowest emissive ones. These are limiting srSMLM because the final image resolution depends on the accuracy of each individual localization measurement (the higher the number of emitted photons, the better the pointing precision) and on the spatial density of emitters localized in the final image (Nyquist sampling theory). Some recent attempts have been made for improving srSMLM resolutions either by improving the photon detection efficiency (9), proposing new fluorophores (20) or by enabling higher emitter densities (22, 23) - but the improvements remain modest. In this work, we aimed to advance srSMLM through data post-processing. Recent developments in deep learning methods now offer powerful analysis tools that have the potential to outperform conventional image processing (24, 25). For example, content-aware image restoration (CARE) (26) can improve and restore the quality of under-exposed or under-sampled images and makes it possible to recover important biological information from noisy images. CARE network architectures are based on convolutional U-nets (27). These deep artificial convolutional network architectures can be trained end-to-end with relatively few images and with relative quick training time to denoise and restore images.

**Fig. 1.**
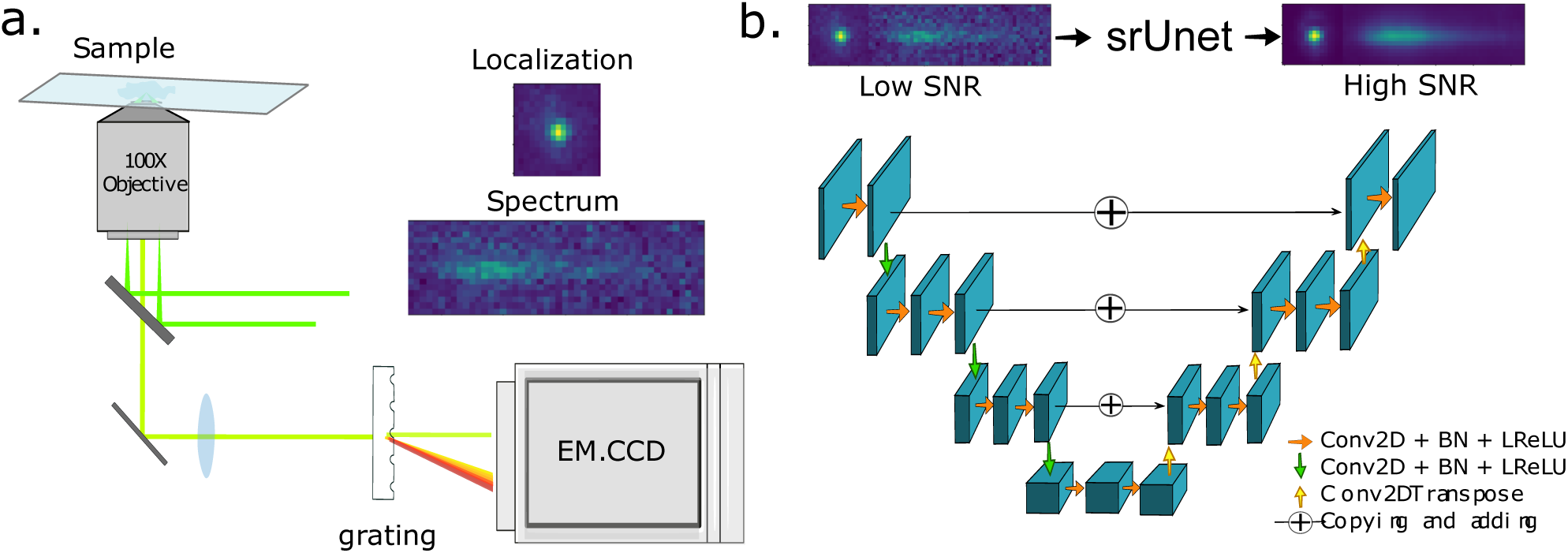
srSMLM and srUnet principles. **a. Schematic representation of srSMLM**. In srSMLM, the diffraction-limited image (PSF) of a single emitter is recorded in a spatial space - allowing its localization to be estimated with high accuracy. Simultaneously, part of the emission light is diffracted and the emission spectrum is recorded in a spectral space on the same detector - giving access to spectroscopic information on the emitter. **b. srUnet architecture**. srUnet is based on a Unet architecture. A Unet is a convolutional neural network made of a contracting path (left part of the U - operating 2D convolutions (Conv2D), batch normalization (BN) and using Leaky Rectified Linear Activation (LReLU) as activation function) and a symmetric expanding path (right part of the U: based on Transposed convolution (Conv2DTranspose) layers also called deconvolution layers) with cross connections between some same-sized parts of the down-sampling and of the up-sampling paths (horizontal arrows). srUnet can be trained end-to-end to restore high signal-to-noise ratio (SNR) from low-SNR image.

We presented here srUnet, a U-net-based image processing for srSMLM that enhances the spectral and spatial signals and compensates for the photon loss inherent in recording the spectral component.

## Results

### Convolutional network

Convolutional neural networks are very efficient at semantic segmentation, classification, image denoising, domain translation, or reconstruction (28, 29). U-net are particular convolutional networks made of a contracting path (left hand side of the U) and a symmetric expanding path (right hand part.) (Fig. 1 b.) with cross connections between same sized parts of the down-sampling and the up-sampling paths (27, 30). In image denoising, U-nets can efficiently restore high signal to noise ratio (SNR) images from low SNR images. The model has first to be trained with pairs of images, using the low SNR images as inputs and their corresponding high SNR images as targets. Once trained, the network can be used to predict high SNR images from any new low SNR images with good generalizing properties for images unseen during the training.

#### Generating a training data set

We first built a training data set made of noisy low-SNR images (x_train) paired with their corresponding high-SNR images (y_train).

Because of the stochastic nature of the emissions in SMLM it is improbable to match low and high SNR acquisition sequences composed of exactly similar localizations in space and time at the scale of a field of view. To circumvent this, we worked on 64×16 pixels image patches - themselves a concatenation of a 16×16 pixels container centered around the PSF in the spatial channel and a 48×16 pixels container collecting the corresponding spectra in the spectral channel (Fig. 2 a. and b.). At the localization scale, collecting the same single molecule several time with different SNR is possible.

**Fig. 2.**
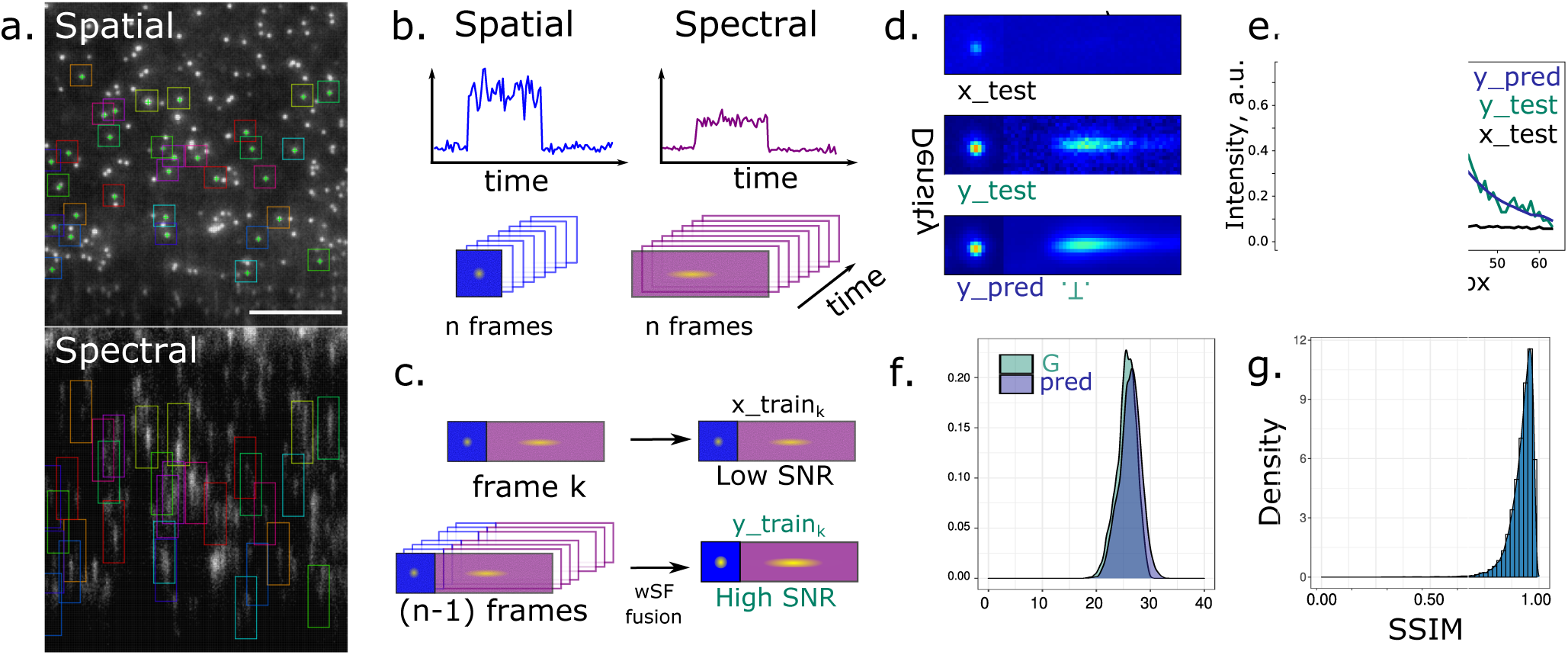
Training data set and performance of the srUnet to learn to predict ground-truth images. **a. Raw srSMLM image (scale bar 10 µm)**. The PSF of a single emitter in the spatial channel can be associated to its spectral decomposition in the spectral channel. The total signal of a single emitter is detected in two distinct domains. **b. Cluster of “mergeable” frames of a single molecule**. In single molecule microscopy, fluorescent molecules can be detected in their ‘on’ state for more than one frame. Molecules appearing on several consecutive frames are usually merged as a single localization. **c. Building srUnet training data-set**. Here we used signal appearing on consecutive frames to generate the srUnet training data-set. All but one image of a cluster were fused using a weighting based on their spatial frequencies to generate a reference image with higher signal to noise ratio (ground truth image or y_train). The image left is used as a train low SNR image (x_train). The training set was composed of *k* ≈ 265, 000 pairs of images and split in a train and a test data-set. **d. Representative images and e. signal profiles of the test data set and of their srUnet predictions**. The y_pred image predicted by the srUnet for the x_test image appears very similar to the y_test reference image used as a target. **f. Peak signal-to-noise ratio (PSNR) and g. structural similarity index (SSIM)**. The restored images predicted by the network (pred) exhibited an average improvement of ~26 dB, a value very similar to the improvement between the raw images and their corresponding ground truth (G.T.). Similarly, the SSIM values between predicted images and their ground-truth target were very close to one, showing that srUnet was able to predict ground truth quality images from raw noisy data

We imaged the ATTO 542 dye of a DNA-PAINT origami nanoruler (GATTA-PAINT 80RG with, GattaQuant GmbH) immobilized on a microscope glass slide. Each DNA-PAINT origami nanoruler has three docking sites for labelled imaging DNA strands (imagers), creating three aligned localization sites with robust and reproducible mark-to-mark distances of 80 nm (31). To fully control the matching between high and low SNR localizations, we took advantage of the fact that the signal from an emitter can be observed on several consecutive frames during the acquisition - in particular at low laser illumination intensity - generating repeated measurements of the same emitter at a given position (Fig. 2 b.). These repeated measurements can be clustered in space and time and clusters with at least three localizations observed in a radius of less than 10 nm were kept. Within a cluster *c* composed of *n* frames, a random frame *k* was selected to append the training set (x_train_*c*_) while the (*n −* 1) left images of the cluster were fused using Spatial Frequency weighted averaging (wSF fusion) (32) followed by a white top-hat transformation to create the corresponding high SNR image (y_train_*c*_). (Fig. 2 c.). This original approach allowed us to build a training data set made of pairs of low SNR / high SNR images of the same PSF. The training set was composed of pairs of images acquired at different laser intensities to collect different levels of signal and/or noise. We also augmented the training set with images in which the spectral container box was randomly shifted along the spectra axis - mimicking a shift of the emission spectra of the emitter. Finally we introduced about 15% of images without any localization in the training data set to force the network to make parsimonious predictions. Taken together, we generated a training matrix of dimensions (265000, 16, 64, 2) (Fig. 2 c.) that we split in a train (90%) and a test (10%) data set.

#### Training the network

The srUnet was trained on the ~ 240,000 pairs of images of the training set. The training time was about 60 min (200 epochs of 15-20 seconds each) on a GPU (Nvidia GTX 1080 Ti) using keras data generator (33). The ~ 25, 000 pairs of images of the test data set were used to evaluate the quality of the training. Restored images (y_pred) were generated from the x_test images and compared to the y_test images used as a ground truth (G.T.) (Fig. 2 d. and e.). Visually, predicted images were very similar to the targeted G.T. images. As compared to the raw images, the restored images predicted by the network (*Ŷ* or y_pred) exhibited an average improvement of 26 [22-29] dB (mean [CI_95%_]) measured by peak signal-to-noise ratio (PSNR) (Fig. 2 f.). The corresponding value between raw images and their corresponding ground truth was 25.5 [21-29] dB (mean [CI_95%_]). As expected from these very similar PSNR and from the visual impression, the structural similarity index (SSIM), a perceptual metric used to quantify the difference between the y_pred images and their corresponding ground truth images, was 0.93 [0.90-0.96] (median [IQR]) (Fig. 2 g.). These high SSIM values clearly demonstrate that the network was able to predict ground truth quality images from raw noisy data.

#### Exploring the reconstruction performance of the srUnet

To evaluate further the reconstruction performance of the trained network (SSIM evaluated differences at the pixel level), we applied it to the previously unseen images of a validation set built independently from the testing set. We first explored the ability of the srUnet to reconstruct properly images from raw noisy images. We explored whether we could fit a PSF in the reconstructed image if a localization was initially truly present in the G.T. image. From the confusion matrix (Fig. 3 a.) we calculated a sensitivity of 99.5%. The specificity was also very good (Sp= 0.988%). Together, the accuracy reached 99.4%, showing that discrepancies relative to the G.T. data was minimal. To explore the properties of the reconstructed PSF, we fitted the G.T. PSF and the reconstructed PSF of the validation data set using the same software (Peakfit) with same setup parameters. Compared to the G.T. PSF, we observed a small systematic bias both on the center position of the 2D-Gaussian fits, with about 4 nm and about 2 nm on the X and Y axes, respectively (Fig. 3 b.), The standard deviation (SD) distributions of the circular 2D-Gaussian fits were very similar (Fig. 3 c.). The presence of the limited spatial bias is hard to interpret because of the unknown true position of the emitting molecules in the sample. This systematic bias is not critical as it only results in a translation of a few nanometers of the reconstructed image. The number of detected photons by emitters calculated as the integral of the 2D-Gaussian fits were greatly increased in the reconstructed srUnet image as compared to the the original raw image. The number of photons increased by a factor of 4 to 15, with a clear tendency for the least emissive emitters to be more improved than the most emissive ones (Fig. 3 d. - black dots). Interestingly, the number of photons in the reconstructed images were very similar the the number of photons retrieved in the G.T. ones, showing that the srUnet was able to learn to reconstruct images with G.T. qualities (Fig. 3 d. - green dots).

**Fig. 3.**
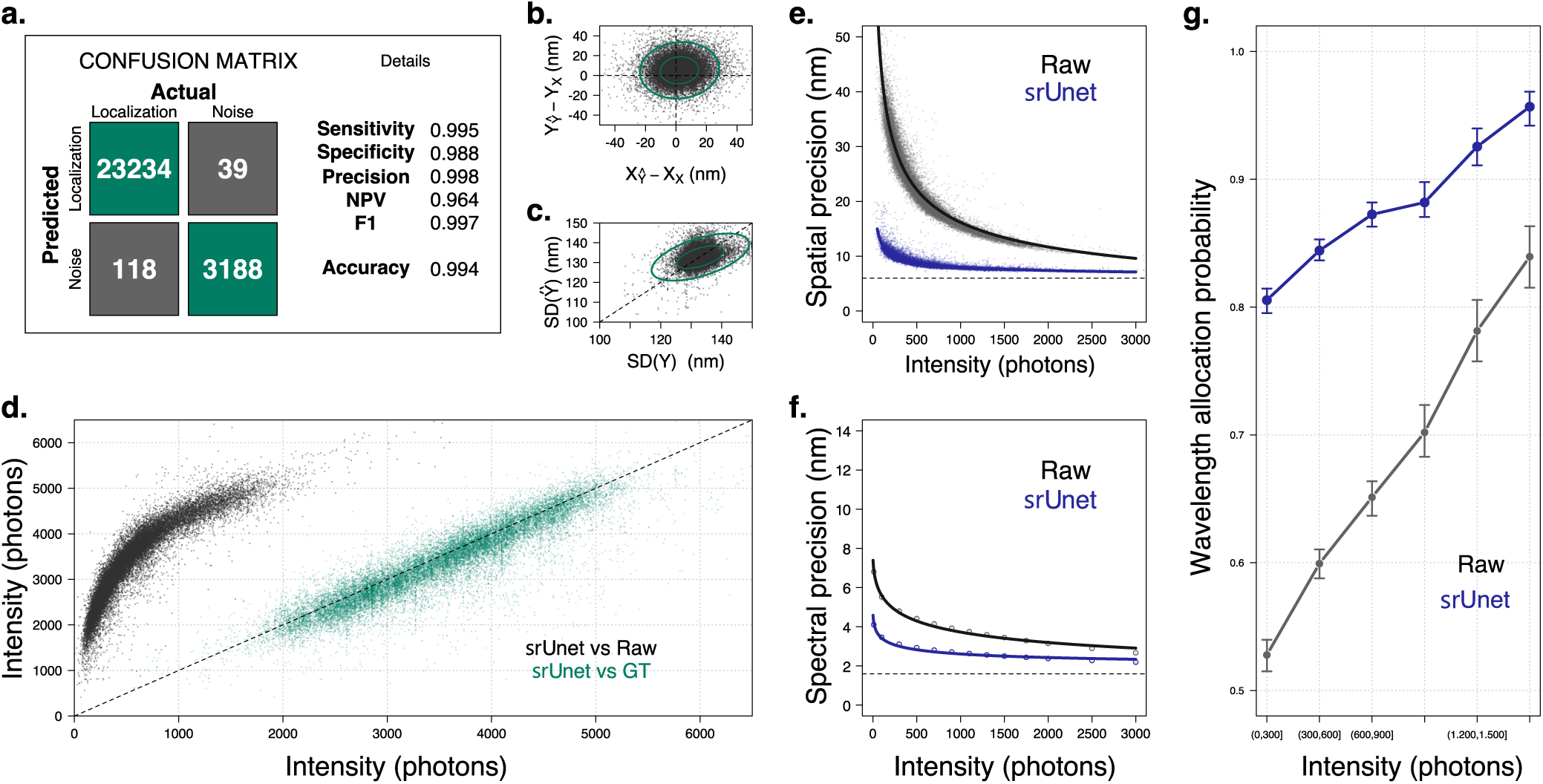
U-net performance evaluated on a validation set. **a. Confusion matrix and localization detection performance metrics**. The localizations retrieved on srUnet restored images were compared to the actual localizations. **b. Center position bias (in nm) and c. standard deviation change of the 2-D Gaussian fits of the PSF**. A few nanometer systematic bias was found in the localization coordinates of the srUnet images using the raw data as a reference, but was not observed in the standard deviation of the 2D Gaussian fits. **d. Comparison of the number of collected photons between the srUnet and (black) raw images or (green) ground truth images**. Interestingly, the improvement factor provided by the srUnet was increasing with the decrease of the raw signals. **e. Spatial and f. spectral precision as a function of the number of collected photons**. We found an ultimate spatial precision of about 6 nm and an ultimate spectral precision of about 1.9 nm. The use of srUnet resulted in great improvement in both spatial and spectral precision. **g. Wavelength allocation probability as a function of the collected intensity signals**. The fraction of localization for which a wavelength can be assigned increases with the signal intensity of single molecules. The fraction retrieved using srUnet was largely increased as compared to raw data and was at least as good as the one obtained for the highest signals in the raw data.

Then we explored comparatively the spatial and spectral precisions achievable as a function of the number of photons. In the spatial domain, typical photons values obtained during srSMLM (400-1,500 photons) led to a spatial pointing precision of 25 nm to 13 nm in the raw images (Fig. 3 e.). The ultimate spatial precision of the instrument (that is with an infinite photon number) was 6 ± 0.5 nm. In line with the enhanced signal in the reconstructed images, the precision calculated from the reconstruction of the same images with the srUnet was much better and in the range of 9 nm to 7nm. For the intensity signals ranging from 400 to 1,500 photons, the spatial precision remained close to the ultimate achievable value of about 6 nm. Interestingly, the spatial precision of the srUnet images was improved over the entire intensity range. The precisions of lowest emitting molecules with the srUnet were at least as good as those obtained for the most emissive molecules in the raw data. A similar trend was observed for the spectral precision. In Fig. 3 f., the conditional standard deviation of the retrieved wavelengths were calculated as the root square of the squared distance from the mean distance (the residuals) of a regression modeling the wavelength as a function of the signal intensity. The ultimate spectral precision was found to be about 1.9 ± 0.3 nm. At the typical intensity of 1,000 photons, the spatial precision was 2.6 ± 0.1 nm for the srUnet images as compared to 3.9 ± 0.2 nm for the raw data.

Finally, we quantified the fraction of localizations for which the spectral parameters were retrieved as a function of their intensityies In the raw data, and as expected, the wavelength allocation probability increased with the intensity, reaching about 80% for the highest emissive molecules and only about 50% for the lowest intense signals. Compared to the raw data, the srUnet wavelength allocation probability was largely improved with a minimal value of about 80% at the lowest intensity to nearly 95% for the highest intensities.

All these results showed that the localization and the spectral precisions of low-emitting species were greatly improved by the srUnet. Their precisions remained as good as those obtained with a high photons budget in the raw data, even for emitting species whose fluorescence intensity was reduced up to 10 times as compared to the best acquisition conditions. The fraction of localizations whose signal is both spatially and spectrally interpretable was also significantly improved and was well above the usual fraction retrieved in the absence of srUnet. Taken together, the use of srUnet resulted in improved pointing precisions and increased the density of spots characterized both spatially and spectrally. As these two parameters define the quality of super-resolution images, srUnet advances Spectrally-Resolved Single Molecule Localization Microscopy to reach performances close to classical SMLM.

#### Effect of spectral shift on the srUnet images

srSMLM has been elegantly used for single-molecule polarity detection (6, 18–20) where local hydrophobicity is probed by slight spectral shifts of spectrally responsive (solvatochromic) fluorophores. It was therefore important to explore the ability of the network to reconstruct images in the presence of spectral shifts of the emission signals. To do so, we artificially added a defined spectral shift to the validation images by shifting the position of the container in the spectral domain relative to the position of the localization container (Fig. 4 a.). We randomly reduce the distance between the containers to induce spectral shifts of 6, 12 or 18 pixels (Fig. 4 b.). The wavelengths at the maximum of the emission spectra were then determined in the raw and in the srUnet images. The distributions of wavelengths at the maximum of the emission spectra are shown in Fig. 4 b.. As expected, the centers of the distributions were shifted to the red with the increase of the added shifts, both in the raw images and in the srUnet images. Distributions retrieved in the raw images appeared wider than the ones of the srUnet images - in line with the fact that uncertainty on the spectra positions are reduced with srUnet. Plotting the mean retrieved wavelengths at the maximum of emission as a function of the added shift (Fig 4 c.) showed that the srUnet preserves the linearity of the response as demonstrated by *r*^2^ values of 0.993 for the srUnet and *r*^2^ = 0.978 for the raw images. In full line with the calibration curve used to characterize the spectral component of srSMLM data (6), we found a slope of 2.1 ± 0.1 nm per pixels for the spectral resolution of the setup.

**Fig. 4.**
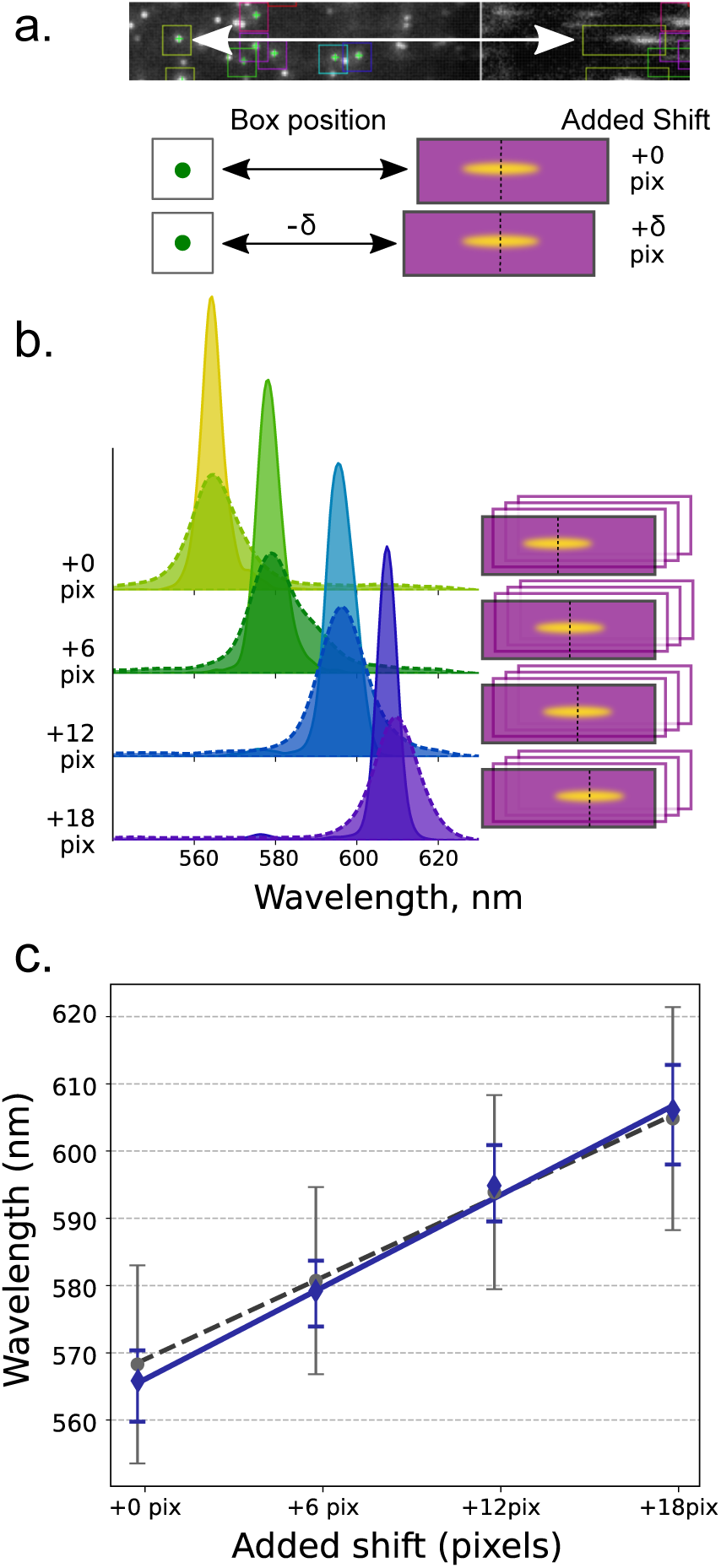
Image reconstruction in the presence of a spectral shift. **a. Synthetic spectral shift**. An artificial shift in the emission spectra is added by shifting the position of the container in the spectral domain relative to the position of the localization container. **b. Retrieved distributions of the wavelength at the maximum of the emission spectra**. Distributions measured in raw data (dashed lines) were systematically wider as compared to those measured in srUnet images (continuous lines). **c. Mean wavelength at the maximum of emission as a function of the added shift**. srUnet preserves the linearity of the response (*r*^2^ = 0.993).

These data clearly demonstrate that srUnet can be applied for single-molecule polarity sensing experiments as srUnet preserves the linearity of the experimental response (when using a transmission grating for light dispersion). This property is also important for multicolour experiments.

#### Multicolour imaging with single laser excitation and fluorophores with overlapping spectra

Successful applications of srSMLM in multicolour imaging largely rely on its ability to achieve true color imaging (5, 8, 9) - offering the potential to clearly outperform multicolour measurements based on the detection of signals in spectrally separated channels. The ability to record the full emission spectra makes srSMLM suitable for multicolour imaging with closely emitting fluorophores, limiting chromatic aberrations and allowing single laser excitation. Even though the complete spectra fluorophores are captured in srSMLM, the fluorophores distinctions are generally made on the basis of their spectral weighted means or positions of their emission maximum - exposing to misidentifications. Improving spectral signals and precision is expected to result in more easily identifiable spectra.

We imaged small fixed and permeabilized Gram negative rod-shaped bacteria with two fluorophores in PAINT mode (3). We used Nile-Red, a solvatochromic dye labelling preferentially lipid membranes and emitting in this sample in the range of 590-620 nm based on local hydrophobicity (34), and POPO-III iodide - a symmetric cyanine dye dimer with high affinity for DNA, poorly emitting in water but showing strong signal when bound to DNA. POPO-III is emitting at its maximum at ~ 575 nm. This is to our knowledge the first reported use of POPO-III as a PAINT probe. Both dyes are excited by a solid state 532 nm laser source and imaged in TIRF mode, limiting the excitation in the sample to the bacterial membrane and nucleoid close to the glass-slide surface. As seen in Fig. 5 a., the experimental emission spectra of POPO-III and Nile Red were largely overlapping. The effect of srUnet on improving the quality of the two-color srSMLM images is double. First, the number of localizations with characterized wavelength information is largely increased, resulting in higher localization densities in the super-resolution images (Fig. 5 b,c,d *vs* e,f,g). Then, the improved precision of the spectral dimension resulted in easier fluorophore determination. In this sample, spatial cross-talk between the two fluorophores is expected because in TIRF mode the Nile Red signals localized in the membrane are projected partially in the same areas as the POPO-III signals staining the DNA rich nucleoid of the bacteria. As a result, the two fluorophores are not spatially separated, challenging fluorophore identification. A better fluorophore discrimination is obtained using U-net. This was easily observed on the transverse section showing better contrasts between the expected Nile Red stained membrane and the POPO-III labelled nucleoid (DNA rich area) (Fig. 5 h-i). Additionally, the distribution of the wavelengths at the maximum of emission of the Nile Red was very similar if the dye was present alone or in mixture with POPO-III - whatever the order of addition of the two fluorophores in the sample. (Supplementary fig Sx) Together, we demonstrate that srUnet improves multicolour imaging in a demanding small sample where the labelled components are not spatially exclusive - even using fluorophores with overlapping spectra and excited by a single laser.

**Fig. 5.**
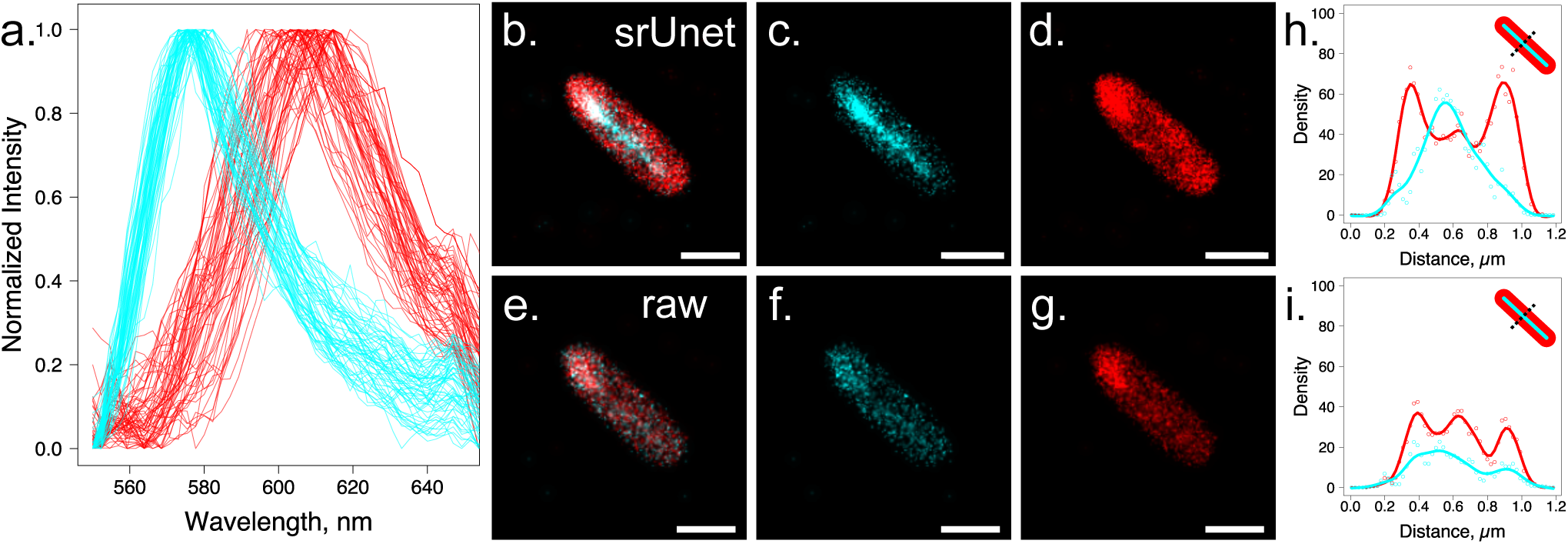
Multicolour srSMLM in bacteria with Nile Red and POPO-III fluorophores. **a. srUnet augmented experimental spectra**. About 200 experimental normalized spectra of single molecules of POPO-III (cyan) and Nile Red (red) are represented to illustrate the spectral overlap between the two fluorophores. **b.-g. srSMLM images of fixed *Pseudomonas aeruginosa***. (scale bar 1 µm) (b. and e.) For visualization, true colours where changed for cyan (*λ ≤* 580 nm) or red (*λ >* 585 nm). Reconstruction are represented using (b.) or not (e.) srUnet. The localizations were also split to represent the reconstruction of the spatial positions of POPO-III (c. and f.) and Nile Red (d. and g.) **h.and i. Transverse intensity profiles of the localizations density in the image in b. (h.) or e. (i.)**, showing better contrasts between the Nile Red stained membranes and the POPO-III labelled DNA-rich nucleoid using srUnet.

## Conclusions

In this work, we trained a deep convolutional U-net (srUnet) to restore in post-processing high signal-to-noise (SNR) images from srSMLM acquisitions. This was made possible by building an original training data set based on spatial-frequency weighted image fusion of the same PSF recorded over several frames. We showed that localization and spectral precision of low-emitting species remain as good as those obtained with a high photons budget, even for emitting species whose fluorescence intensity is reduced up to 10 times. The spatial precision of srSMLM localizations obtained with srUnet are very similar to those obtained in classical SMLM. The use of srUnet therefore almost fully compensates the photon loss inherent to the recording of the spectral information. As a consequence of the signal restoration, the fraction of localizations whose signal is both spatially and spectrally interpretable was largely improved and well above 80% under all tested conditions. srUnet was also able to reconstruct properly images in the presence of spectral shifts. Finally, we demonstrated that srUnet is easily applicable to real data collected in biological samples and greatly facilitates multicolour imaging with single laser excitation - even for fluorophores with overlapping emission. spectra As the trained srUnet showed good generalization properties for Nile Red and POPO-III dyes, our approach is expected to work for many other fluorophores and other technical setups designed for srSMLM.

If the contributions of the network are extremely interesting as is, it could be envisioned to further advance the approach. This might include coupling the inference on the localization coordinates and spectral wavelengths directly within the deep learning network. A deep network could also be built to directly learn image reconstruction from raw data and derived versions could also be created for multicolour spectrally-resolved single molecule tracking experiments.

srSMLM can greatly benefit from the latest improvements in deep learning. Coupling srSMLM with deep learning could overcome the potential actual limitations of srSMLM and make its use as broad and easy as SMLM.

## Material and Methods

### Spectrally-resolved Single molecule localization microscopy

SMLM imaging was performed on a home-built bespoke Micro-Manager controlled Olympus IX-81 inverted optical microscope equipped with a 100X 1.4 NA oil-immersion objective (Olympus - Japan) and a z-drift control and auto-focus system (ZDC Olympus). A 532 nm continuous wavelength diode-pumped solid-state laser (Coherent Sapphire SF) was used as a light source. The fluorescent emission was collected through the same objective using a 532 nm dichroic mirror (Di02-R532 - Semrock) and filtered by a longpass filter (532 LP Edge Basic, Semrock). A physical aperture (VA100/M, Thorlabs) and a transmission diffraction grating (300 Grooves/mm 8.6° Blaze Angle - GT13-03 Thorlabs) were mounted before the detector to create the spatial and spectral channels on a 512 × 512 pixels electron multiplied charged coupled device (EMCCD) camera (ImagEM - Hamamatsu Photonics - Japan).

Typical image stacks ranging from 10,000 to 15,000 frames with exposure time 40 ms were recorded. Image were processed by PeakFit (part of GDSC SMLM2 plugin) and/or ThunderSTORM an open source software packages for Fiji in order to retrieve molecules localization coordinates. The procedure described in Bongiovanni et al. (6) was used for spectral calibration. All subsequent and additional image processing were performed using self-written scripts in Python (version 3.8).

### DNA origami nanorulers

Green (ATTO 542) nanorulers with mark-to-mark distances in the sizes 80 nm (Gatta-Paint) were available commercially (GATTAquant GmbH, Germany) and used according to the manufacturer recommendations.

#### Culture and sample preparation of bacteria

Briefly, *Pseudomonas aeruginosa* (PAO1 - DSM 22644) were grown overnight in 5 mL lysogeny broth (LB) (L3152, Sigma Aldrich) at 30 °C under 220 rpm orbital shaking. Cells were then diluted at OD_600nm_ = 0.1 and grown for an additional 2 to 3 hours. Cell cultures were pelleted by centrifugation, washed three times with Phosphate Buffered Saline (PBS) before being fixed using para-formaldehyde (PFA 4%) for 15 minutes at room temperature under gentle agitation. Cells were finally washed three times with PBS, permeabilized with lysosyme and stored at +4 °C. The detailed protocol can be found elsewhere (35).

#### Preparation of the coverslips for PAO1 imaging

Glass coverslips (0.13 mm thickness, 20 × 20 mm)(Knittel Glass) were cleaned using an argon plasma cleaner (PDC-002, Harrick Plasma) for 20 min. Frame-seal slide chambers (9 × 9 mm, Bio-rad, USA) were affixed to the glass coverslips. The chamber was filled with poly-L-lysine solution (0.01% w/v, P4707 Sigma Aldrich) to fully coat the coverglass surface, incubated for at least 10 min and then washed ones with PBS buffer. Fifty µL of fixed PAO1 cell solution were allowed to settle on PLL treated coverslips for 10 min before being washed ones with PBS buffer. The slide was transferred to the microscope stage using an Attofluor cell chamber (A-7816 Thermofisher) and optically coupled to the objective lens through index-matching immersion oil (n=1.518, Olympus, Japan). Diluted solution of Nile Red (Invitrogen) or POPO-III-iodide (Invitrogen) dyes were added directly on the sample on the microscope stage.

### srUnet

The srUnet was adapted from the princeps Unet paper (27) and built using Keras library package (2.4.0) running on Tensorflow backend (2.5.0). The model was composed of 6 × 6 convolutions with Exponential Linear Unit (ELU) activation function followed by batch normalization and a Leaky Rectified Linear Unit (Leaky ReLU) (Fig. 1, b). The model had about 1.3 · 10^6^ trainable parameters. The root mean squared error loss function was minimized using RMSprop optimizer. The model was trained using batch size of 70 over about 200 epochs. The training time was fast and the algorithm converged smoothly.

#### Weighted image fusion

Image fusion was used to gather all the important information from multiple images. We used spatial frequency (SF) measures to weight the different images as SF measures the overall information level in an image. SF was defined as

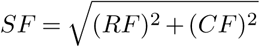

where RF and CF are respectively the row frequency

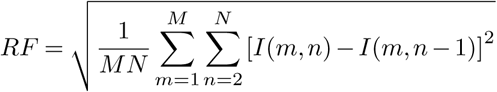

and column frequency

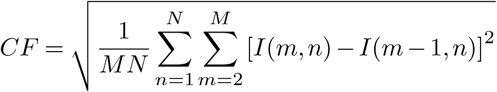

in which M and N are the number of pixels of the (MxN) grey image and I(m,n) is the intensity of the pixel of coordinates (m,n).

#### Metrics for classification performance and Image quality assessment

We used Accuracy (*Acc* = (*tp* + *tn*)*/N*, Specificity (*Sp* = *tn/*(*tn* + *fp*)), Sensitivity or Recall (*Se* = *tp/*(*tp* + *fn*), Positive Predictive Value or Precision (*Precision* = *tp/*(*tp* + *fp*), Negative Predictive Value (*NPV* = *tn/*(*fn* + *tn*)) and F_1_ (*F*_1_ = 2.(*precision*.*recall*)*/*(*precision* + *recall*) metrics for performance evaluation. All these measurements were expressed in terms of *tp* = true positive, *tn* = true negative, *fp* = false positive and *fn* = false negative.

Image quality assessment were reported using classical MSE, PSNB and SSIM metrics (36) defined as:

Mean Squared Error (MSE)

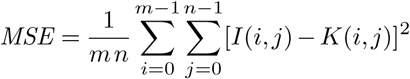

Peak signal-to-noise ratio (PSNR) in dB

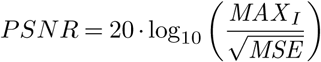

with *MAX*_*I*_ = 2^16^

Structural similarity index measure (SSIM)

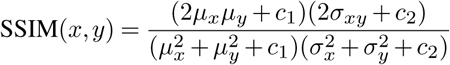

## Software and Hardware Availability

The code used for the training set generation and training the network is available on https://github.com/hmanko/srSMLM_DeepL.

Most figures of this article can be reproduced independently using the code found on this repository.

## ACKNOWLEDGEMENTS

We thank Dr Isablle Schalk for providing PAO1 bacteria strain.

This research was supported by the ITI Innovec (IdEx (ANR-10-IDEX-0002) and SFRI (ANR-20-SFRI-0012)). HM was funded by a 3-years contract (Contrat doctoral) from Ministère de la Recherche, France. YM is grateful to the Institut Universitaire de France (IUF) for support and providing additional time to be dedicated to research.

## AUTHOR CONTRIBUTIONS

These contributions follow the Contributor Roles Taxonomy guidelines: https://casrai.org/credit/. Conceptualization: H.M., J.G; Data acquisition and curation: H.M., J.G.; Formal analysis: H.M.,J.G.; Funding acquisition: J.G., Y.M.; Investigation: H.M., J.G.; Methodology: H.M., J.G.; Project administration: J.G.; Resources: J.G., Y.M.; Software: H.M., J.G.; Supervision: J.G.; Validation: J.G.; Visualization: H.M., J.G.; Writing of original draft: H.M., J.G.; Writing review & editing: all authors.

## COMPETING FINANCIAL INTERESTS

The funders had no role in study design, data collection and interpretation, or the decision to submit the work for publication. The authors declare no other competing financial interests.

## Notes

### Competing Interest Statement

The authors have declared no competing interest.

